# Age-associated Differences in the Human Lung Extracellular Matrix

**DOI:** 10.1101/2022.06.16.496465

**Authors:** M.L. Koloko Ngassie, M. De Vries, T. Borghuis, W. Timens, Don D. Sin, D. Nickle, P. Joubert, P. Horvatovich, G. Marko-Varga, J.J. Teske, J.M. Vonk, R. Gosens, Y.S. Prakash, J.K. Burgess, C.A. Brandsma

**Affiliations:** University of Groningen, University Medical Center Groningen, Department of Pathology and Medical Biology, Groningen, Netherlands; University of Groningen, University Medical Center Groningen, Groningen Research Institute for Asthma and COPD, Groningen, Netherlands; University of Groningen, University Medical Center Groningen, Department of Epidemiology, Groningen, Netherlands; Centre for Heart Lung Innovation at St. Paul’s Hospital, University of British Columbia, Vancouver, Canada; Gossamer Bio, San Diego, CA, USA; Institut Universitaire de Cardiologie et de Pneumologie de Québec, QC, Canada; University of Groningen, Department of Analytical Biochemistry, Groningen Research Institute of Pharmacy, Groningen, Netherlands; Lund University, Center of Excellence in Biological and Medical Mass Spectrometry, Biomedical Center, Lund, Sweden; Department of Anesthesiology and Perioperative Medicine, Mayo Clinic, Rochester, Minnesota, USA; University of Groningen, Department of Molecular Pharmacology, Groningen, Netherlands

**Keywords:** Aging, extracellular matrix, lung, remodelling, airway wall, blood vessel, parenchyma, bronchial epithelium, collagen

## Abstract

**Introduction:** Extracellular matrix (ECM) remodelling has been associated with chronic lung diseases. However, information about specific age-associated differences in lung ECM is currently limited. In this study we aimed to identify and localize age-associated ECM differences in human lung using comprehensive transcriptomic, proteomic and immunohistochemical analyses.

**Methods:** Our previously identified age-associated gene expression signature of the lung was re-analysed limiting it to an aging signature based on 270 control patients (37-80 years) and focused on the Matrisome core geneset using geneset enrichment analysis. To validate the age-associated transcriptomic differences on protein level, we compared the age-associated ECM genes (F <0.05) with a profile of age-associated proteins identified from a lung tissue proteomics dataset from 9 control patients (49-76 years) (FDR<0.05). Extensive immunohistochemical analysis was used to localize the age-associated ECM differences in lung tissues from control patients (9-82 years).

**Results:** Comparative analysis of transcriptomic and proteomic data identified 7 ECM proteins with higher expression with age at both gene and protein level: COL1A1, COL6A1, COL6A2, COL14A1, FBLN2, LTBP4 and LUM. With immunohistochemistry we demonstrated higher protein expression with age for COL6A2 in whole tissue, parenchyma, airway wall and blood vessel, for COL14A1 in bronchial epithelium and blood vessel, and for FBLN2 and COL1A1 in lung parenchyma.

**Conclusion:** Our study revealed that higher age is associated with lung ECM remodelling, with specific differences occurring in defined regions within the lung. These differences may affect lung structure and physiology with aging and as such may increase susceptibility for developing chronic lung diseases.

**Key messages:** *What is already known on this topic:* summarise the state of scientific knowledge on this subject before you did your study and why this study needed to be done. ❖ In animal models, it has been demonstrated that aging alters the composition of the lung ECM, with more deposition of collagen and degradation of elastin. Similar ECM differences have been observed in age-associated chronic lung diseases, including COPD; moreover, we observed in lung tissue that several ECM genes associate differently with age in COPD patients compared to non-COPD controls(1). Detailed knowledge on age-associated changes in specific ECM proteins as well as regional differences within the lung is lacking.

*What this study adds:* summarise what we now know as a result of this study that we did not know before. ❖ We identified 7 age-associated ECM proteins i.e. COL1A1, COL6A1, COL6A2 COL14A1, FBLN2, LTBP4 and LUM with higher transcript and protein levels in human lung tissue with age. Extensive immunohistochemical analysis revealed significant age-associated differences for 3 of these ECM proteins in specific compartments of the lung, with the most notable differences in the blood vessels and parenchyma.

*How this study might affect research, practice, or policy:* ***s**ummarise the implications of this study*. ❖ The identification of age-associated differences in specific human lung ECM proteins lays a new foundation for the investigation of ECM differences in age-associated chronic lung diseases. Additionally, examining the function of these age-associated ECM proteins and their cellular interactions in lung injury and repair responses may provide novel insight in mechanisms underlying chronic lung diseases.

## Introduction

Aging is a natural phenomenon that affects molecular and biological processes and subsequently also the structure and function of tissues and organs including the lung, contributing in the impairment of the tissue and organ homeostasis(2). Lung aging is characterized by structural and physiological differences including larger size of alveoli, less elasticity, thickening of the small airway wall, worse lung function, and remodelling of the extracellular matrix (ECM)(3–5). Significant changes such as decreased elastin and increased collagen deposition have been observed in lung ECM from old mice(6, 7). Several features of aging have been found to be more pronounced in chronic lung diseases including chronic obstructive pulmonary disease (COPD) with a high prevalence in the elderly(3, 8).

Given its role as a provider of the architectural structure, mechanical support and regulator of several biological processes(9), the ECM plays an important role in organ homeostasis. The molecular composition, biological and mechanical properties of the ECM are tissue specific(9). The lung ECM is composed of a complex combination of elastin, collagens, glycosaminoglycans, proteoglycans and glycoproteins. Components of the ECM have specific functions in organs, accordingly elastin provides the lung its extension and recoil properties(10), collagens provide tensile strength and regulate cellular migration and adhesion, glycosaminoglycans regulate growth factor activity and lung viscoelasticity through their hydration properties(9). In our previous study, a gene set enrichment analysis (GSEA) revealed a significant negative enrichment of the ECM-receptor interaction pathway with age in COPD compared to non-COPD tissue, indicating that ECM genes change differently with increasing age in COPD patients compared to controls(1). In addition, a recent publication highlighted the positive enrichment of genes of the ECM pathway with age in human lung(11), and Burgstaller and colleagues showed age-associated changes in the expression of ECM proteins in mouse lung tissues(12). Altogether, these studies suggest age-related alterations in lung ECM proteins, however, detailed knowledge is still scarce and information on regional differences within the lung is lacking.

In this study we examined the age-associated differences in ECM proteins at both transcriptional and protein levels in human lung tissue derived from patients with normal lung function and no history of chronic lung disease. Additionally, we performed immunohistochemistry to assess age-associated ECM differences in specific lung compartments, i.e. whole lung tissue, lung parenchyma, airway wall, bronchial epithelium, and blood vessels.

## Materials and methods

More detailed description of used methods is available in the online **supplementary file 1.**

### Procurement of lung tissues

The transcriptomic and proteomic data were derived from lung tissues from control patients (For the numbers of patients used see analysis sections below) with normal lung function and no history of chronic lung disease. The characteristics of these patients were previously described by De Vries *et al* and Brandsma *et al*(1, 13). The lung tissues used for immunohistochemistry were derived from patients undergoing therapeutic lung resection surgery for cancer at the University Medical Center Groningen (Groningen, The Netherlands), which were part of the HOLLAND (HistopathOLogy of Lung Aging aNd COPD) project, or the Mayo Clinic, (Rochester, Minnesota, USA). The HOLLAND project is a large immunohistochemistry study performed at the department of Pathology and Medical Biology of the UMCG aiming to identify and correlate differences in several ECM, inflammatory, epithelial and senescence markers in serial sections from the same lung in relation to chronic lung disease (i.e. COPD and IPF) and aging. The control patients from Groningen all had normal lung function, i.e. FEV1/FVC > 70%, no lung function data were available for the patients from Mayo Clinic. Patients were non-smokers, ex-smokers, or current smokers, with an age range between 9 and 82 years.

### Transcriptomic analyses

We re-analysed our previously identified age-associated gene expression signature of the lung, limiting it to an aging signature based solely on the 270 non-disease control patients(1). In short, linear regression analysis adjusted for the potential confounders sex, smoking status (ex or current smokers) and technical variation using principal components explaining >1% of the variation was performed using R software. The three cohorts Groningen, Laval and Quebec were analysed separately and combined by a meta-analysis. To correct for multiple testing, the Benjamini-Hochberg false discovery rate (FDR) was applied. Next, we determined the enrichment of ECM-associated genes using Gene Set Enrichment Analysis (GSEA 4.1.0), including all genes present in the Matrisome geneset (NAB A_ Matrisome; M5889, Molecular Signatures Database v7.4). All Matrisome genes with a significant enrichment score were defined as our age-associated ECM genes.

### Proteomic analyses

We assessed the association between age and protein expression in 9 non-disease control patients present in our previously published lung tissue proteomics dataset(13). The edge R package version 4.1.0 was used for linear regression of the data following a negative binomial distribution and using edgeR as described in Brandsma et al but using only age as factor involved in the regression analysis(13). FDR p<0.05 was considered significant.

### Immunohistochemical staining

Immunohistochemical staining for collagen type I alpha 1 (COL1A1) COL6A1, COL6A2, COL14A1, fibulin-2 (FBLN2), latent transforming growth factor beta binding protein 4 (LTBP4) and lumican (LUM) was performed in paraffin-embedded lung tissue derived from 64 control patients (Figure 1A).

**Figure 1:**
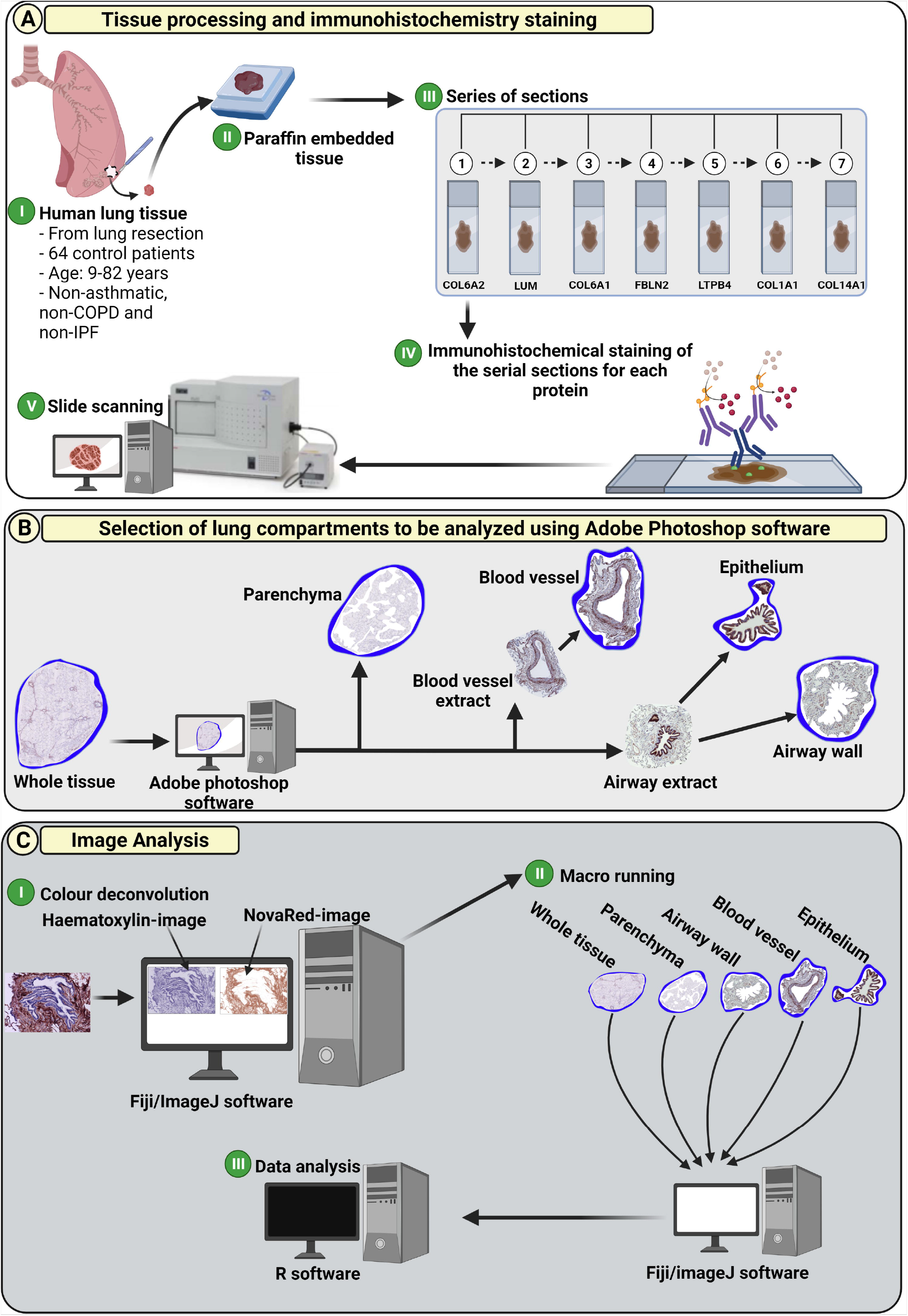
Tissue processing, immunohistochemistry, and image analysis. **(A)** Lung tissues were embedded in paraffin and cut in serial sections. The sections were then immunohistochemically stained for ECM proteins including, COL1A1, COL6A1, COL6A2, COL14A1, FBLN2, LTBP4 and LUM. After the staining, the images were captured using the Hamamatsu NanoZoomer 2.0HT digital slide scanner at magnification of 40x. (**B**) The digital images were checked for their quality and once validated the whole tissue, parenchyma, airway wall, airway epithelium and blood vessel wall areas were isolated and cleaned (artifacts were removed). (**C**) For the image analysis, a macro was developed to separate the haematoxylin and NovaRed image components using the colour deconvolution plugin in ImageJ. Afterwards, a specific developed macro was run for each staining, followed by data analysis using R software. Figure was created with BioRender.com.

### Image analyses

Different compartments of the lung including whole lung tissue, parenchyma, airway wall, bronchial epithelium, and blood vessel (both arteries and veins) were analysed separately. Specific regions were captured using Aperio ImageScope software V.12.4.3 (Leica Biosystems, Nussloch, Germany) and Adobe Photoshop software (Adobe Inc. California, United States) (Figure 1B). Fiji/ImageJ software (14) was used to quantify the mean intensity and percentage of positive stained area for each protein (Figure 1C). Data analyses were performed using R software V.4.0.0 (Boston, Massachusetts, USA) using the following formula’s:

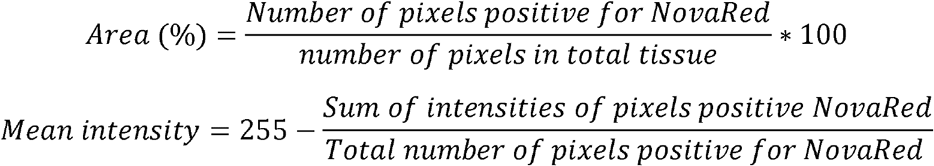

### Statistical analysis

Linear regression and linear mixed model analyses with a random effect on intercept, correcting for sex and smoking status, were performed to determine the association between age and each protein using SPSS software V.27 (IBM Corp. in Armonk, NY, USA). P<0.05 was considered significant.

## Results

### Patient Characteristics

The clinical characteristics (Table S1) of the control patients included in the transcriptomics dataset included 192 ex-smokers and 78 current smokers with a normal lung function and an age range of 37-80 years. The clinical characteristics (Table S2) of the proteomics dataset included 9 ex-smoking controls with normal lung function and an age range of 49-76 years.

The clinical characteristics of the 64 control patients used for IHC staining are summarized in Table 1. 44 lung tissues were collected in Groningen and included 16 never-smokers, 18 ex-smokers and 10 current smokers. The 20 lung tissues collected in Rochester were all derived from never smokers.

**Table 1:** Patient characteristics

|  | Groningen | Rochester |
| --- | --- | --- |
| **Number, n** | 44 | 20 |
| **Age, years (range)** | 9-82 | 18-80 |
| **Male/female, n** | 16/28 | 5/15 |
| **Never-smoker, n** | 16 | 20 |
| **Ex-smoker, n** | 18 | 0 |
| **Current-smoker, n** | 10 | 0 |
| **FEV<sub>1</sub>, mean +/- SEM** | 101 +/-15.46 | NA |
| **FEV<sub>1</sub>/FVC ratio, mean+/-SEM** | 0.76 +/- 0.05 | NA |

The FEV<sub>1</sub> and FVC data were not available for lung tissues from Rochester and for few cases from Groningen. FEV<sub>1</sub>: forced expiratory volume in 1 s, FVC: forced vital capacity, NA: not applicable

### Gene expression signature for lung aging in non-disease controls

To determine the association between age and ECM gene expression, we first assessed the age-associated gene expression differences in the non-disease control lung tissues. In total 4201 probes corresponding to 4147 unique genes were significantly associated with age; of which 2247 probes coding for 2226 genes showed a higher and 1954 probes coding for 1939 genes a lower gene expression with higher age (full list in **supplementary file 2**). The top 10 most significantly higher and lower expressed genes are depicted in Table 2.

**Table 2:** The top 10 genes with higher and lower expression in relation to Age in non-disease control lung tissue

| Genes higher expressed with higher age |  |  |  | Genes lower expressed with higher age |  |  |  |
| --- | --- | --- | --- | --- | --- | --- | --- |
| Probe number | Gene name | Meta summary | Adjust BH | Probe number | Gene name | Meta summary | Adjust BH |
| 26059 | COL16A1 | 0.012 | 1.08E-15 | 44483 | GGCT | -0.007 | 4.26E-10 |
| 27901 | EDA2R | 0.025 | 1.02E-14 | 27529 | KIAA1543 | -0.010 | 2.72E-09 |
| 11226 | GALNT6 | 0.013 | 2.63E-14 | 43729 | ZNF518B | -0.010 | 4.18E-09 |
| 23264 | MXRA8 | 0.001 | 1.38E-13 | 26110 |  | -0.012 | 1.93E-07 |
| 26591 | LOXL1 | 0.016 | 1.35E-12 | 1257 |  | -0.006 | 3.48E-07 |
| 46136 | NFASC | 0.014 | 5.93E-12 | 5658 | FANCE | -0.009 | 9.38E-07 |
| 36412 |  | 0.013 | 8.85E-12 | 30969 | TMEM41B | -0.006 | 1.98E-06 |
| 18268 | ITGBL1 | 0.022 | 1.41E-11 | 4655 | EFNA1 | -0.009 | 2.97E-06 |
| 17176 | MMP2 | 0.011 | 2.15E-11 | 44483 | GGCT | -0.012 | 3.41E-06 |
| 16116 | F8 | 0.023 | 4.69E-11 | 27529 | KIAA1543 | -0.007 | 4.70E-06 |

### Enrichment of Matrisome pathway among the age-associated gene signature in the lung tissue

As we were specifically interested in age-related differences in ECM gene expression, we next assessed the enrichment of the ECM genes among the ranked gene list using GSEA analysis for the Matrisome geneset consisting of 1026 ECM (-associated) genes of which 915 were present in our data.

The GSEA analysis showed a strong, significant positive enrichment of the Matrisome pathway ((enrichment score of 0.39) Figure 2) with a total of 318 core enriched ECM genes. The top 25 most significant core enriched ECM (-associated) genes are shown in Table 3 (full list in **supplementary file 3**).

**Figure 2:**
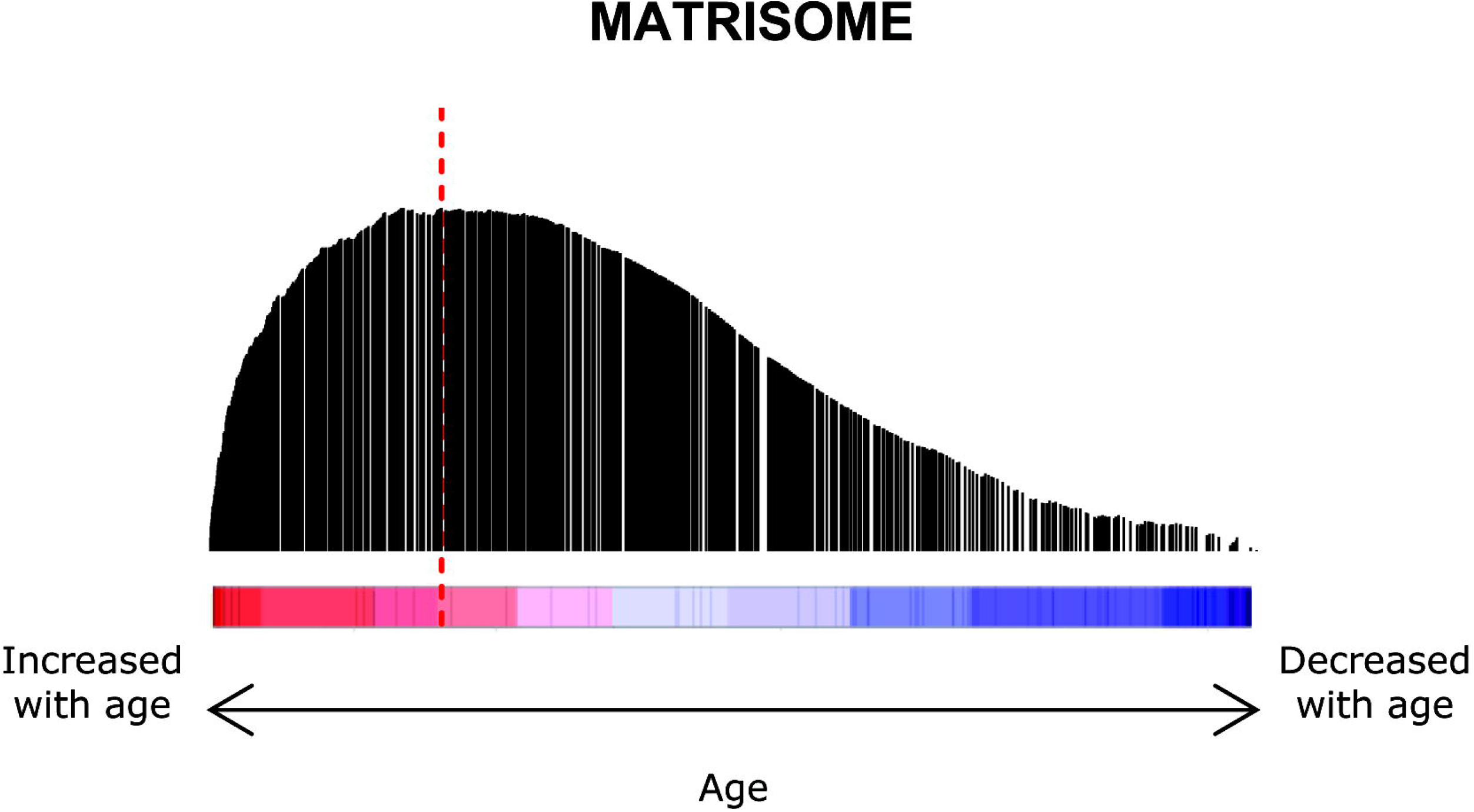
Enrichment of Matrisome pathway among the age-associated gene signature in non-disease control lung tissue. The vertical bar indicates the position and the heigh indicates the enrichment score of a specific gene of the Matrisome. Genes are ranked from the highest to the lowest expressed (from left to right) with increasing age. The red dotted line indicates the highest enrichment score (0.39) of the Matrisome with higher age.

**Table 3:**
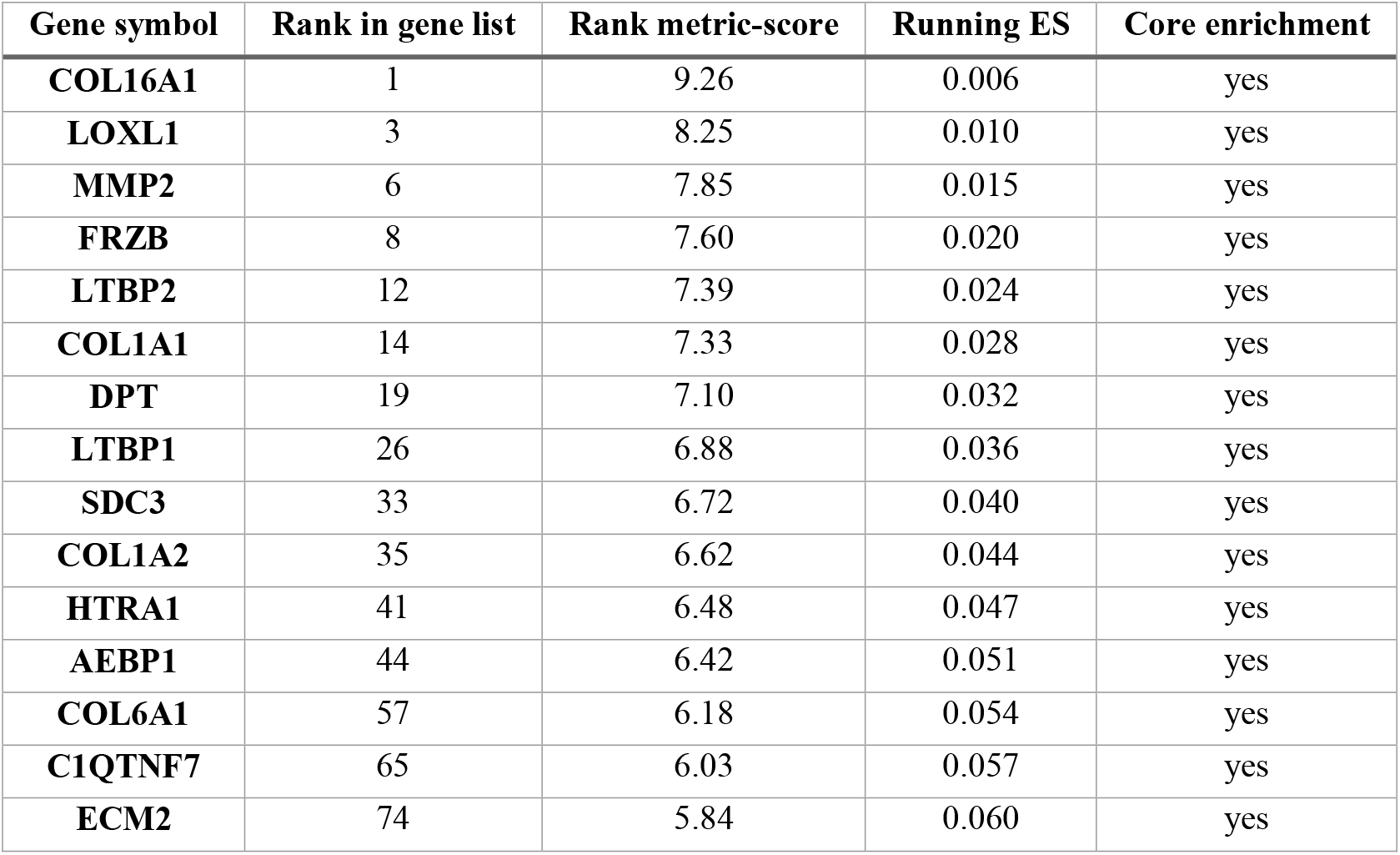
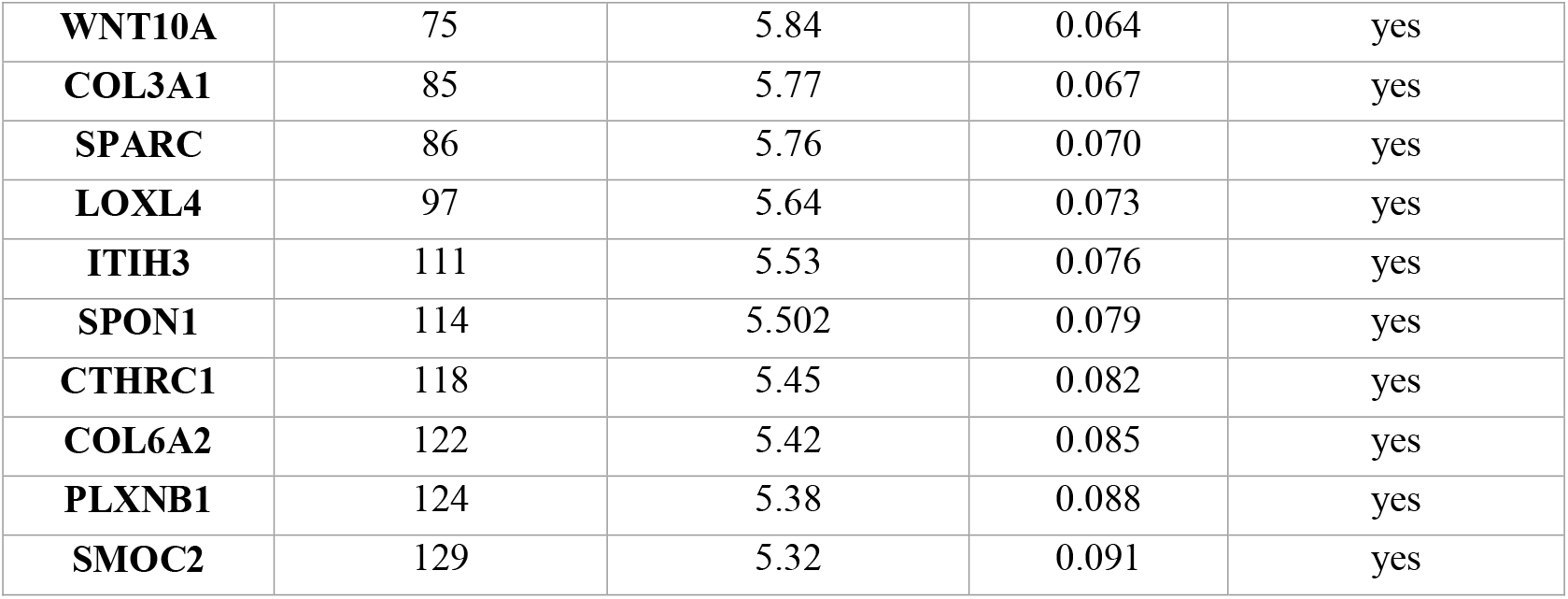
GSEA analysis revealed the association of ECM gene expression in non-disease control lung tissue with aging. The top 25 ranked enriched ECM genes. ES; enrichment score.

### ECM proteins are among the top ranked age-associated proteins in lung tissue

Next, we determined the association between age and protein levels in the lung in the proteomic dataset. We identified 25 differentially expressed proteins, including 20 proteins of which levels were higher with higher age and 5 proteins of which the levels were lower with higher age (Figure 3). Among the 20 proteins with higher protein levels with higher age were several ECM proteins including COL1A1, COL6A1, COL6A2, COL14A1, FBLN2, LTBP4 and LUM.

**Figure 3:**
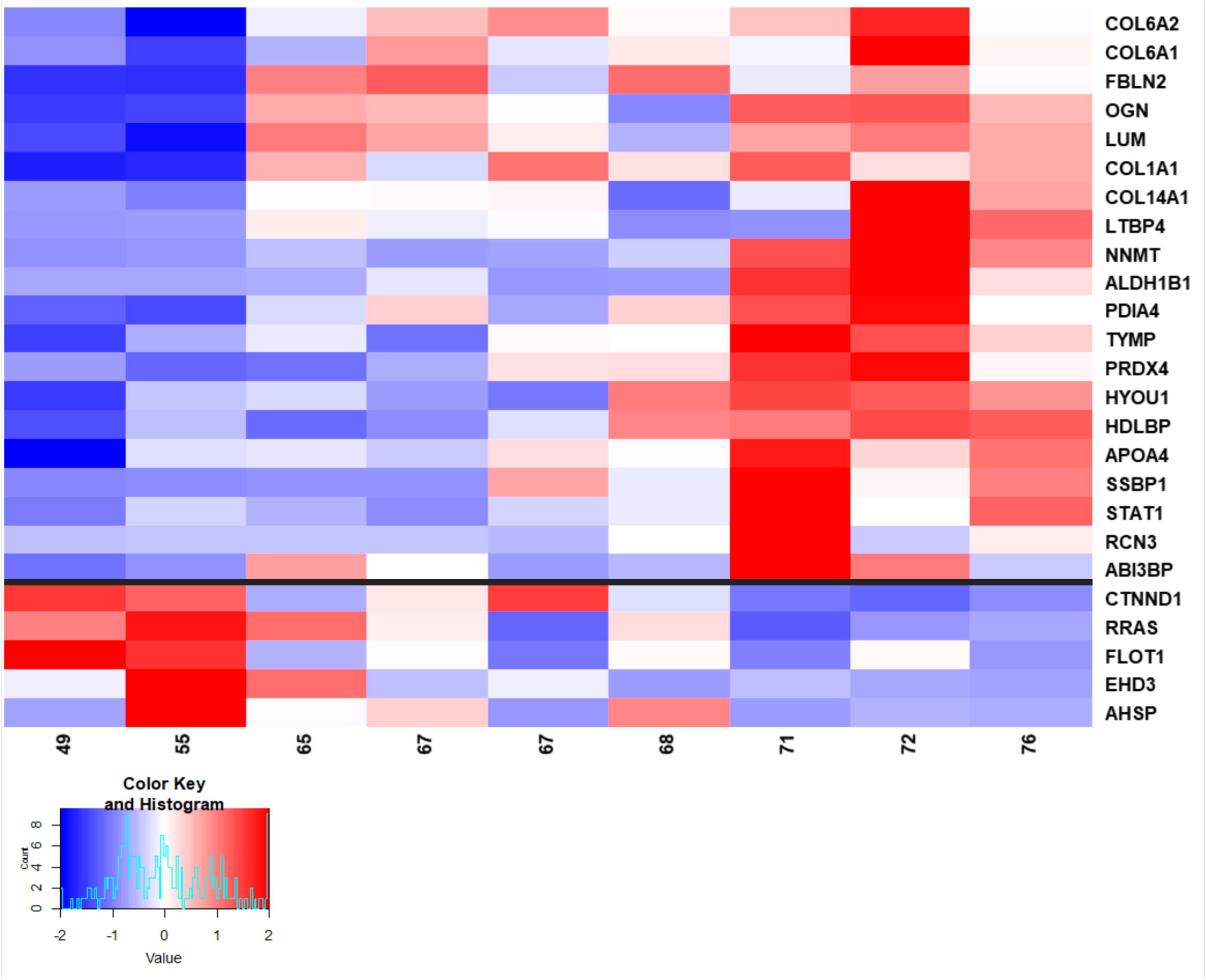
Heatmap of age-associated proteins in the lung of non-disease control patients. The heat-map shows the results of the proteomic analysis of human lung tissue from control patients with normal lung function. The upper part of the map shows the significantly upregulated and the lower part the significantly downregulated proteins with age. For the statistical analysis, p-values were corrected using the Benjamini-Hochberg (FDR) method using a threshold of 0.05. FDR: fold discovery rate.

### Overlap in age-associated ECM proteins in lung tissue on transcript and protein level

Next, we determined the overlap between the age-associated ECM genes identified in the transcriptomic analysis and the age-associated proteins in the proteomic analysis. We identified 7 ECM proteins that were significantly associated with age with correlation in the same direction in both datasets, i.e. higher levels with higher age. This included COL1A1, COL6A1, COL6A2, COL14A1, FBLN2, LTBP4 and LUM (Figure 4A). The beta values for the age association of the 7 age-associated ECM genes from the transcriptomics analysis are shown in the forest plot. (Figure 4B). The 7 overlapping age-associated ECM proteins were among the top 8 proteins in the proteomics analysis (Figure 3).

**Figure 4:**
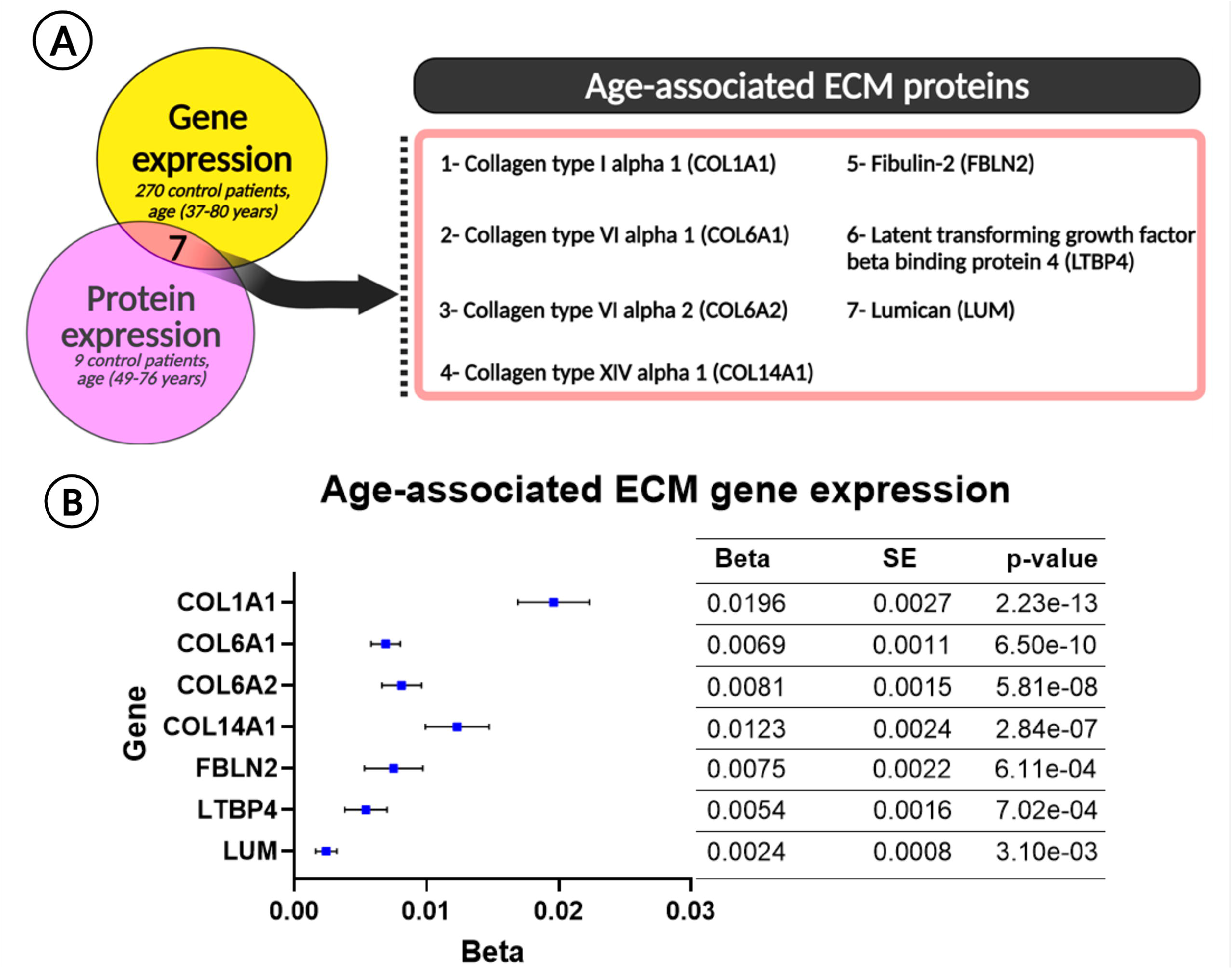
Overlap in age-associated ECM proteins in the lung tissue on transcript and protein level. **(A)**transcriptomic and proteomic results were examined for the overlapping of ECM gene and its encoded protein. Seven ECM proteins including COL1A1, COL6A1 COL6A2, COL14A1, FBLN2, LTBP4 and LUM were significantly higher with age on gene and protein level**. (B)**The beta-coefficients for age of the 7 ECM genes overlapping with encoded ECM proteins are shown in the forest plot. For the statistical analysis, p-values were corrected using the Benjamini-Hochberg (FDR) method using a threshold of 0.05. FDR: fold discovery rate. SE: standard error.

### Localization of age-associated ECM proteins in lung tissue

The immunohistochemically stained tissues were assessed for the localization of the age-associated ECM proteins in the lung (Figure 5). All ECM proteins including COL1A1, COL6A1, COL6A2, COL14A1, FBLN2, LTBP4 and LUM showed positive staining in the parenchyma. The collagens COL1A1, COL6A1 and COL6A2, as well as FBLN2, showed positive staining in the submucosa and adventitia of the airway wall, and in the adventitia of the blood vessel. Additionally, FBLN2 staining was also present in the wall of small vessels. The localization of FBLN2 is suggestive of colocalization with elastin (15). Interestingly, COL14A1 showed positive staining in the airway adventitia, airway smooth muscle (ASM) layer and the apical region of the bronchial epithelium and macrophages. In the blood vessels, COL14A1 was localized in the endothelium and partly in the smooth muscle layer. LTBP4 staining was most prominent in blood vessel smooth muscle and was also present in the bronchial epithelium, ASM layer and macrophages. LUM staining was present in the bronchial epithelium, ASM layer, endothelium and blood vessel smooth muscle layer. An overview of the strenght and localization of the positive stained age-associated ECM proteins in different compartments of the lung is summarized in Table 4.

**Figure 5:**
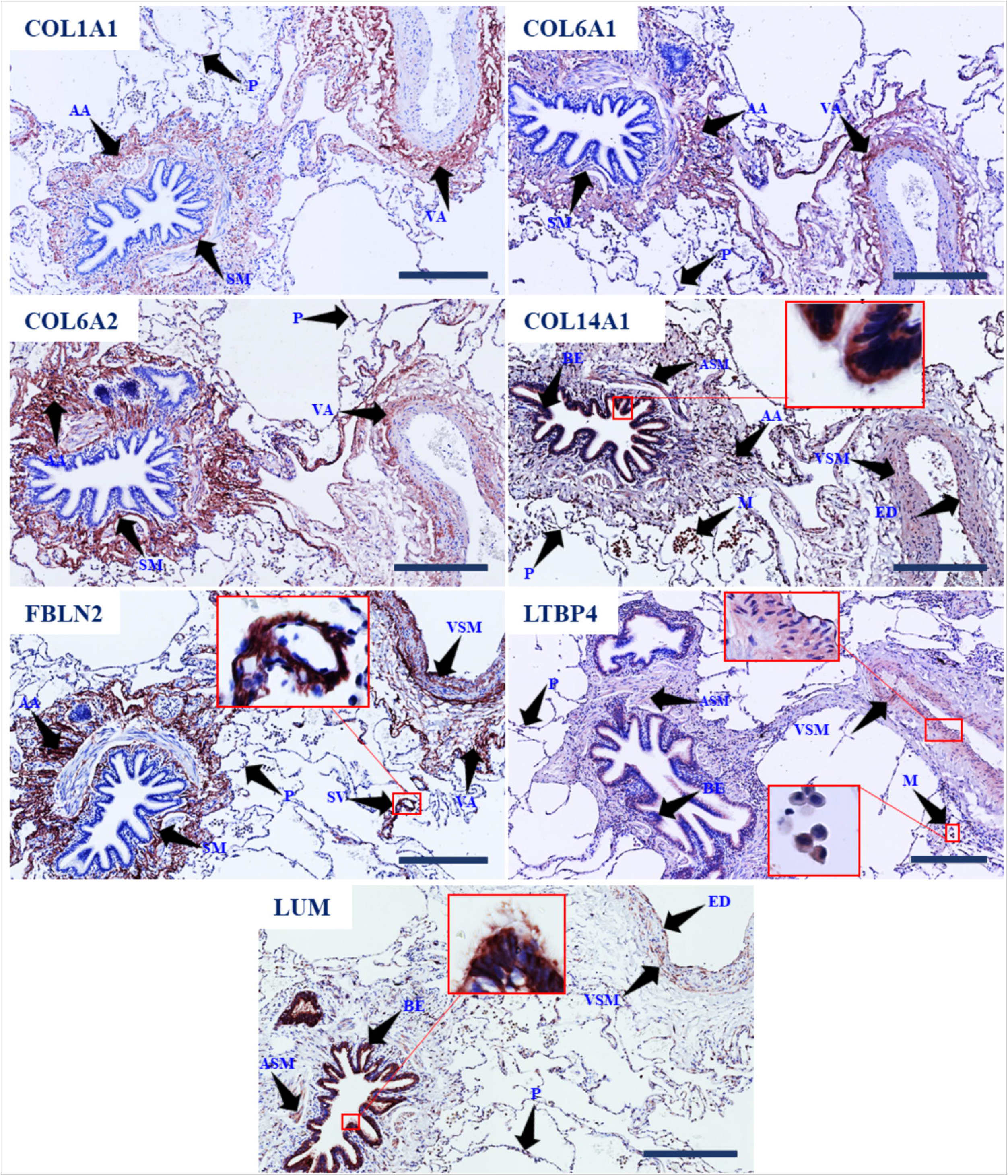
Localization of age-associated ECM proteins in human lung tissues. Following the immunohistochemical staining the age-associated ECM proteins were localized in the lung tissue. All the stained ECM proteins including COL1A1, COL6A1, COL6A2, COL14A1, FBLN2, LTBP4 and LUM were located in the parenchyma. COL1A1, COL6A1 and COL6A2 were localized to the submucosa and airway adventitia and blood vessel adventitia. COL14A1 is localized to the ASM layer, bronchial epithelium, endothelium, blood vessel smooth muscle layer and macrophages. FBLN2 is also present in the submucosa and airway adventitia, smooth muscle layer of large blood vessel, and in the wall of small vessel. LUM and LTBP4 are both present in the bronchial epithelium, airway wall and blood vessel smooth muscle. Macrophages were also positive for LTBP4 staining, whereas LUM was localized to the ASM layer. Arrows indicate areas of positive staining. Scale bar 300 μm. COL1A1: collagen type I alpha 1, COL6A1: collagen type VI alpha 1, COL6A2: collagen type VI alpha 2, COL14A1: collagen type XIV alpha 1, FBLN2: fibulin-2, LTBP4: latent transforming growth factor beta binding protein 4, LUM: lumican, AA: airway adventitia, BA: blood vessel adventitia, ED: endothelium, BE: bronchial epithelium, ASM: airway smooth muscle, VSM: blood vessel smooth muscle, SM: submucosa, SV: small vessel, M: Macrophage, P: parenchyma.

**Table 4:**
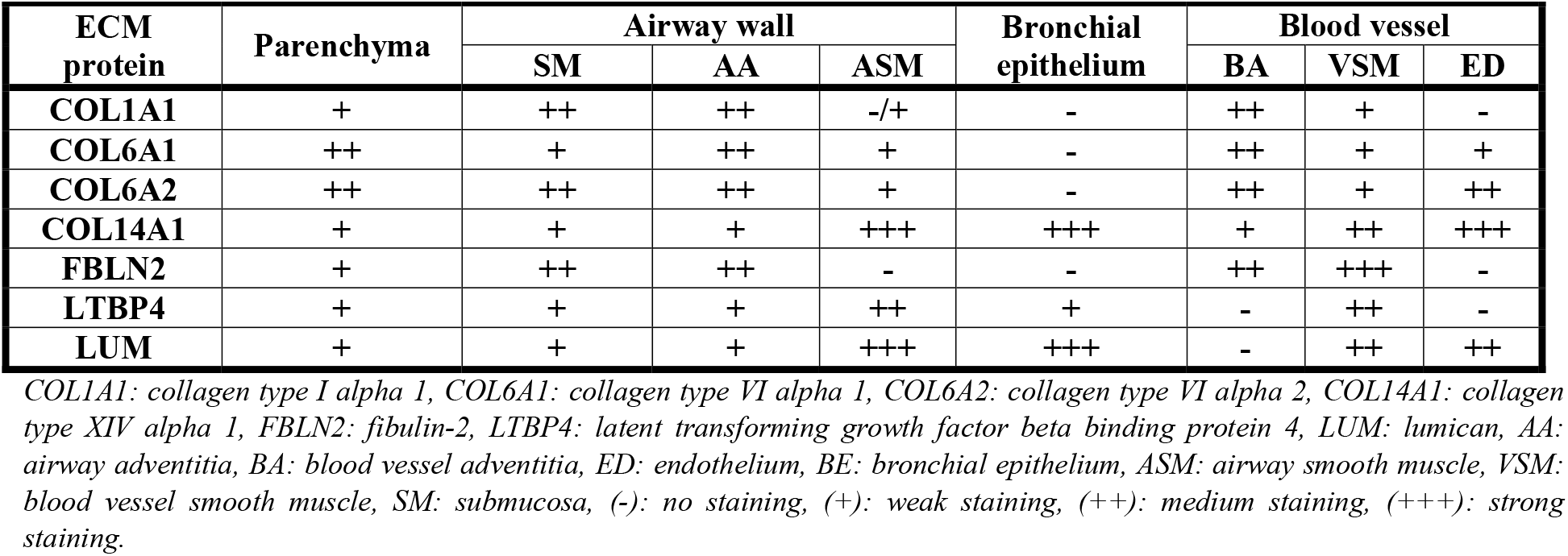
Overview of positive stained areas for age-associated ECM proteins including COL1A1, COL6A1 COL6A2, COL14A1, FBLN2, LTBP4 and LUM and their semi-quantitative score in the lung compartments.

### Evaluation of age-associated ECM protein differences in lung tissue using immunohistochemistry

Following the localization of the 7 age-associated ECM proteins in lung tissue, we determined the age-association in whole lung tissue and in different lung regions, i.e., parenchyma, airway wall, bronchial epithelium and vessel walls with respect to percentage of positive stained tissue area as well as the mean intensitiy of the positive staining.

### Age-association of ECM staining in whole lung tissue

Analysis of the whole tissue showed a significantly positive association between age and the percentage area and the mean intensity of the COL6A2 positive stained tissue (Figure 6A). For FBLN2, we also found a significantly a positive association between age and the percentage area of positive stained tissue (Figure 6A), but not for the mean intensity. For COL1A1, COL6A1, COL14A1, LTBP4 and LUM positive staining in whole lung tissue, no significant association with age was found.

**Figure 6:**
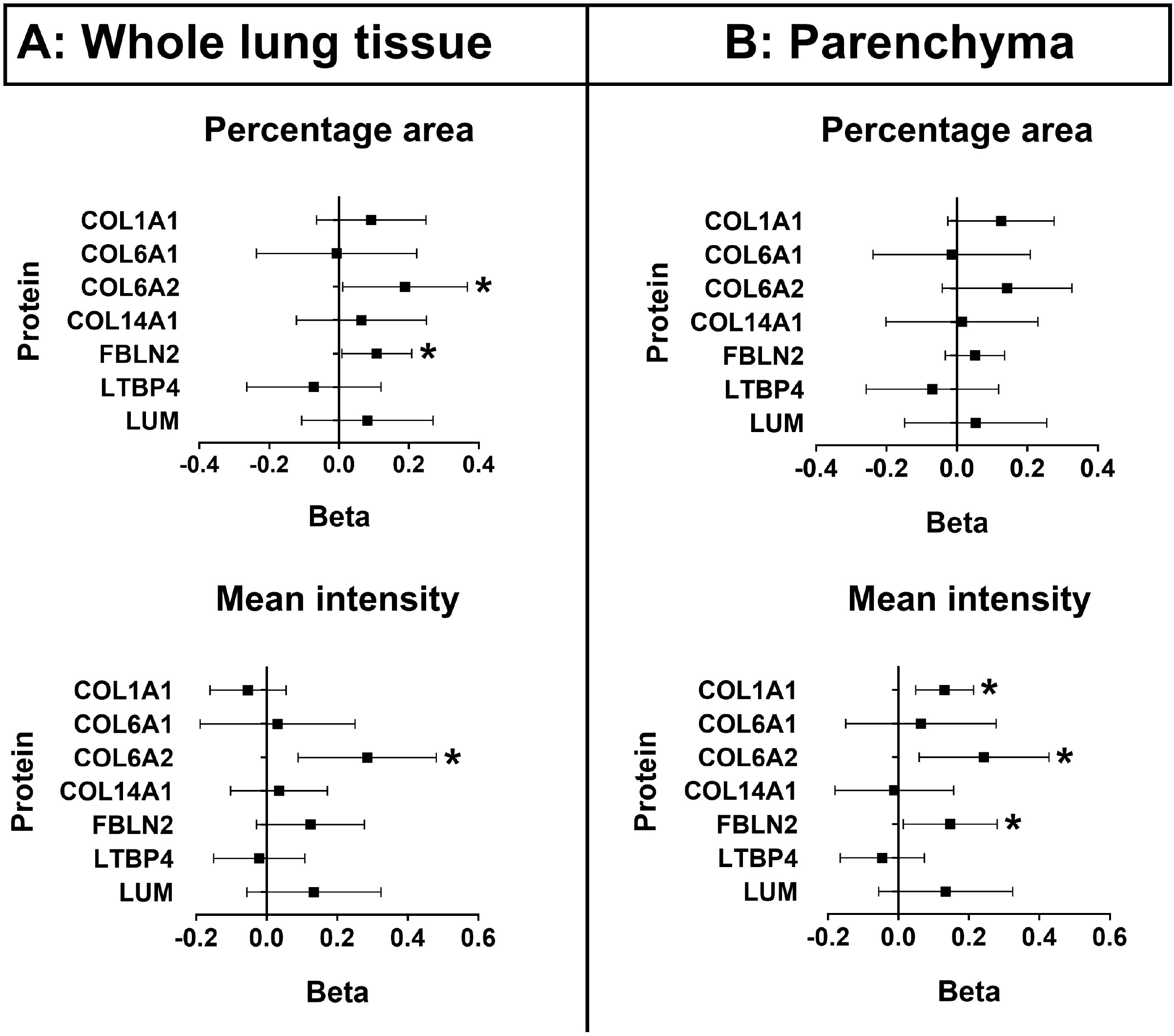
Forest plot of regression estimates for age of the percentage area and mean intensity of age-associated ECM proteins in the whole lung tissue and parenchyma. **(A)** Whole lung tissue sections form control patients without airflow limitation aged from 9-82 were immunohistochemically stained for the 7 age-associated ECM proteins and the positive staining was analysed using Image J software for the percentage area and mean intensity for COL1A1, COL6A1, COL6A2, COL14A1, FBLN2, LTBP4 and LUM positive staining. The analysis of whole lung tissue showed a positive association of percentage area and the mean intensity of COL6A2 positive stained tissue with age. FBLN2 also showed a positive association with age in terms of total positive stained tissue, but no association of the mean intensity of the staining. The percentage area and mean intensity of COL1A1, COL6A1, COL14A1, LTBP4 and LUM positive stained tissue showed no association with age. **(B)** Parenchyma regions isolated from immunohistochemical stained whole lung tissue sections from control patients without airflow limitation aged from 9-82 were analysed using Image J software for the percentage area and mean intensity of COL1A1, COL6A1, COL6A2, COL14A1, FBLN2, LTBP4 and LUM positive staining. The analysis of the parenchyma revealed the positive association of mean intensity of COL1A1, COL6A2 and FBLN2 positive stained tissue with age, but no association was found for the percentage area. Both mean intensity and percentage area of COL6A1, COL14A1, LTBP4 and LUM positive stained tissue showed no association with age. As statistical analysis, the linear regression model adjusted for sex and smoking was performed in SPSS software V.27. The total number of lung tissues used for these analyses varied between 62 and 64. COL1A1: collagen type I alpha 1, COL6A1: collagen type VI alpha 1, COL6A2: collagen type VI alpha 2, COL14A1: collagen type XIV alpha 1, FBLN2: fibulin-2, LTBP4: latent transforming growth factor beta binding protein 4, LUM: lumican. *Significant.

### Age-association of ECM staining in lung parenchyma

The analysis of the parenchymal regions showed a significantly positive association between age and the mean intensity of COL1A1, COL6A2 and FBLN2 positive stained parenchymal tissue (Figure 6B), but no association was found for the percentage area stained. No associations were found between age and COL6A1, COL14A1, LTBP4 and LUM positive stained parenchymal tissue.

### Age-association of FBLN2 and COL6A2 staining in the airway wall

The airway wall was analysed for the percentage area and mean intensity of COL1A1, COL6A1, COL6A2, COL14A1, FBLN2, LTBP4 and LUM positive staining. Age was significantly positively associated with the percentage area of FBLN2 and the mean intensity of COL6A2 in the airway wall. (Figure 7A). However, no age-associated difference was observed for the percentage area of COL1A1, COL6A1 COL6A2, COL14A1, LTBP4 and LUM, and the mean intensity of COL1A1, COL6A1, COL14A1, FBLN2, LTBP4 and LUM positive staining.

**Figure 7:**
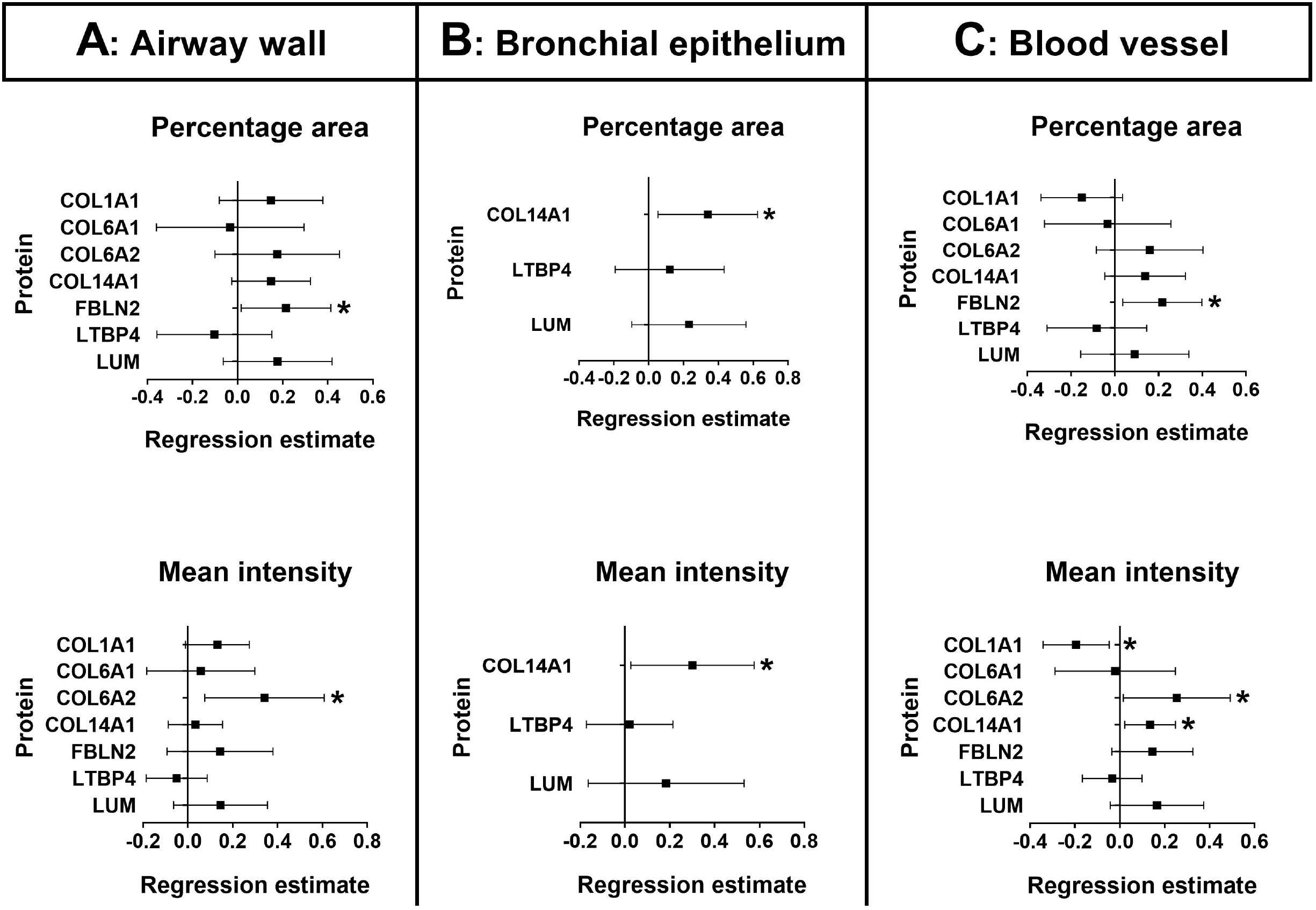
Forest plot of regression estimates for age of the percentage area and mean intensity of age-associated ECM proteins in the airway wall, bronchial epithelium, and blood vessel. Airway wall, bronchial epithelium and blood vessel were isolated from immunohistochemically stained whole lung tissue sections from control patients without airflow limitation aged from 9-82 and analysed using Image J software for the percentage area and mean intensity of COL1A1, COL6A1, COL6A2, COL14A1, FBLN2, LTBP4 and LUM in the airway wall and blood vessel, and COL14A1, LTBP4 and LUM in the bronchial epithelium. **A)** In the airway wall, age was positively associated with the percentage area of FBLN2 and mean intensity of COL6A2. **B)** In the bronchial epithelium, age was positively associated with the percentage area and mean intensity of COL14A1. **C)** The blood vessel showed a positive association between age and the percentage area of FBLN2 and mean intensity of COL6A2 and COL14A1. Age was negatively associated with the mean in intensity of COL1A1. A linear mixed model adjusted for sex and smoking was applied and the regression estimate and 95% confidence intervals for age were represented for each percentage area/mean intensity. The age-associated ECM proteins COL1A1, COL6A1, COL6A2 and FBLN2 were not expressed in the bronchial epithelium. COL1A1: collagen type I alpha 1, COL6A1: collagen type VI alpha 1, COL6A2: collagen type XVI alpha 2, COL14A1: collagen type XIValpha 1, FBLN2: fibulin-2, LTBP4: latent transforming growth factor beta binding protein 4, LUM: lumican, *significant.

### Age-association of COL14A1 staining in the bronchial epithelium

COL14A1, LTBP4 and LUM showed positive staining in the bronchial epithelium and were therefore analysed for their association with age. The percentage area and mean intensity of COL14A1 were positively associated with age (Figure 7B). No age-association was observed for percentage area and mean intensity of LTBP4 and LUM positive staining in the bronchial epithelium.

### Age-association of ECM staining in the blood vessel

The blood vessel wall was analysed for the percentage area and mean intensity of COL1A1, COL6A1, COL6A2, COL14A1, FBLN2 and LTBP4 positive staining. For LUM, the region from tunica media to endothelium of the blood vessel was analysed. FBLN2 was the only age-associated ECM protein that showed a positive association between its percentage area and age. The mean intensity of COL6A2, COL14A1 staining in vessel walls showed a significant positive association with age (Figure 7C). The mean intensity of COL1A1 staining in the blood vessel walls showed a negative association with increasing age. The percentage area of COL1A1, COL6A1, COL6A2, COL14A1, LTBP4 and LUM positive staining; and the mean intensity of COL6A1, FBLN2, LTBP4 and LUM positive staining were not associated with age.

### Age-associated ECM differences are compartment specific

The results of the immunohistochemical analyses are summarized in Figure 8. With respect to the total area of positive ECM staining, COL6A2 was the age-associated ECM protein being higher expressed in all lung compartments including the parenchymal region, the airway wall, and the blood vessel; except in the bronchial epithelium region where it is not expressed. Compartment specific age-associated differences have been observed in the lung for COL1A1, COL14A1 and FBLN2 showing a higher expression in the parenchymal region, bronchial epithelium, and blood vessel wall, respectively. Surprisingly, COL1A1 expression was lower in the blood vessel with increasing age. The blood vessel and the parenchymal regions show the most prominent age-associated differences in their ECM composition.

**Figure 8:**
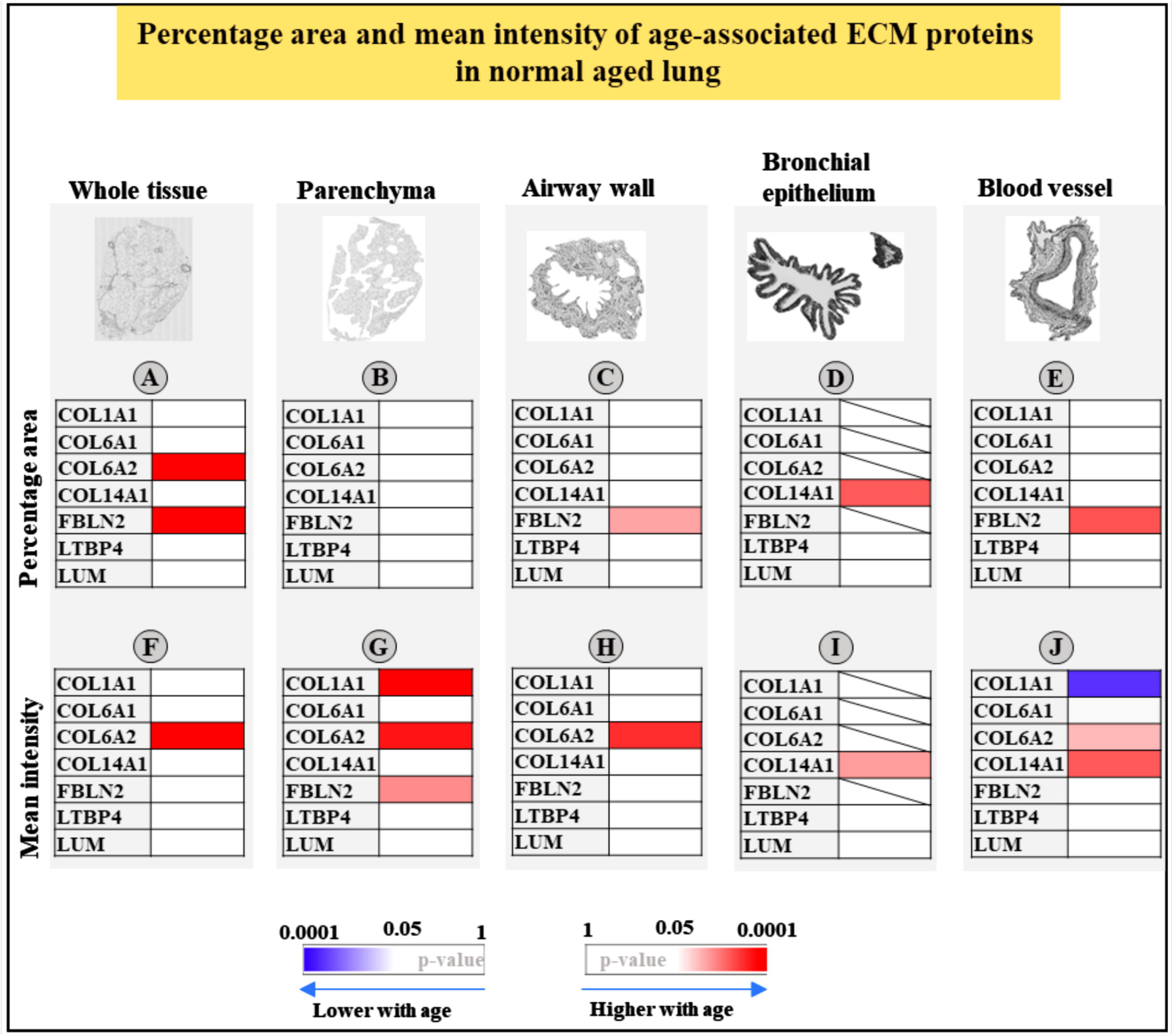
Summary of percentage area and mean intensity of the positive staining of age-associated ECM proteins in different compartments of the lung. In the whole tissue and parenchyma region analysis COL1A1, COL6A2 and FBLN2 show age-associated differences in their percentage area (A and B) and/or mean intensity (F and G), respectively. The percentage area (C) of FBLN2 and the mean intensity (H) of COL6A2 are higher with higher age in the airway wall, while the bronchial epithelium shows a lower percentage area (D) and mean intensity (I) of COL14A1. In the blood vessel, the percentage area of FBLN2 (E) and the mean intensity of COL6A2 and COL14A1 (J) were higher with higher age, while the mean intensity of COL1A1 positive staining was lower with higher age. COL1A1: collagen type I alpha 1, COL6A1: collagen type VI alpha 1, COL6A2: collagen type VI alpha 2, COL14A1: collagen type XIV alpha 1, FBLN2: fibulin-2, LTBP4: latent transforming growth factor beta binding protein 4, LUM: lumican.

## Discussion

Our study describes the age-associated differences in human lung ECM using transcriptomic, proteomic and immunohistochemical analysis. Our results indicate that the human lung ECM remodels with normal aging. The Matrisome pathway, including both ECM and ECM-related proteins, was significantly and positively enriched among the age-associated gene signature in non-diseased control lung tissue. Comparing age-associated transcriptomic and proteomic differences in lung tissue, we identified 7 age-associated ECM (and ECM-associated) proteins being COL1A1, COL6A1, COL6A2, COL14A1, FBLN2, LTBP4 and LUM which all showed higher levels in whole lung tissue with higher age. Subsequent immunohistochemical staining in different compartments of the lung revealed differences in the expression patterns of these ECM proteins with all proteins being expressed in the airway wall and blood vessels and 3 with clear expression in the bronchial epithelium. Age-associated differences were observed for COL6A2 in whole tissue, parenchyma, airway wall and blood vessels, for COL14A1 in bronchial epithelium and blood vessels, and for FBLN2 and COL1A1 in the parenchyma.

Our findings are in line with previous data showing higher expression of collagens and collagen-related proteins in the lung of 24-month-old mice and a higher collagen deposition observed in parenchymal regions of aged mice (16). Other studies specifically showed higher expression of COL1A1, COL6A1 and COL6A2, but a lower expression of COL14A1 with higher age in mouse lung tissue(17, 18).

In our study, COL6A2 was the only protein observed, being higher expressed throughout the different lung compartments. Surprisingly, no age-associated difference was observed for COL6A1 in any of the lung compartments where it was localized. COL6 plays an important role in cell-ECM interaction, through interaction with cell surface receptors including integrins(19). Additionally, COL6 enhances lung epithelial cell spreading and facilitates wound healing(20) and COL6 depletion in mice was linked to altered basement membrane structure and diminished cell-ECM interaction in the lung resulting in an altered pulmonary elasticity and less tolerance for physical exercises(19). These suggest the importance of COL6 in mechanical regulation of the lung, however it remains unresolved why only COL6A2 was higher expressed in our lung tissues and not COL6A1.Thus, it will be interesting to examine the role of COL6A2 in the lung and its implication of lung aging phenomenon.

Higher age is associated with higher expression of COL14A1 in the bronchial epithelium and blood vessel. The role of COL14A1 in collagen fibrillogenesis has been demonstrated, as it restraints collagen fibril diameter(21). Because it is involved in restraining the collagen fibres diameter, COL14A1 possibly involved in the modulation of collagen fibres in the blood vessel wall. COL14A1 is involved in the turnover and differentiation of epithelial cells(17), indicating a role for COL14A1 in the homeostasis of the bronchial epithelium. However, additional investigations are needed for better understanding of COL14A1 in the bronchial epithelium.

Lung elasticity is known to be less with higher age. FBLN2 serves as a bridge between fibrillin and elastin molecules and has shown a strong binding affinity to tropoelastin and COL4(22). Additionally, FBLN2 plays a role in the stabilization of the basement membrane by interacting with COL4(23). More FBLN2 in airway wall and blood vessel wall indicates a possible role in the stabilization of their basement membranes in aged lung. FBLN2 has been identified as a positive regulator of transforming growth factor beta-1 (TGF-ß1) activity(24, 25). FBLN2-KO mice displayed a decreased COL1 expression in the ischemic myocardium(26), which suggests that the increased expression of COL1A1 in the parenchyma in aged tissues maybe linked to increased expression of FBLN2. COL1 contributes to the tissue tensile strength, and its increased deposition may contribute to the lung stiffening with age, as significant increase in stiffness was observed in the parenchyma of old (41-60 year) compared to young (11-30 year) human lung tissues(27). Therefore, more stiffening of the lung with age is associated with the more deposition of FBLN2 and COL1A1. Surprisingly, COL1A1 expression was decreased in aged blood vessel wall, indicating differences in the regulation of COL1A1 expression in the blood vessel compared to the parenchyma and the importance of assessing age-associated differences in different structural compartments in the lung.

The expression of age-associated ECM proteins in different lung compartments may occur through activation of specific biological processes in certain cell types including fibroblasts, myofibroblast and ASM, which are recognized as the principal sources of ECM proteins. An increased proportion of fibroblasts was found in aged human lung compared to the proportion of epithelial cells(4, 28) and our previous work demonstrated a link between cellular senescence and ECM dysregulation in COPD lung fibroblasts. In addition, senescent ASM displayed an higher expression of ECM proteins with higher age(29). These observations suggest that the observed age-associated differences in the lung ECM may be associated with more cellular senescence in aged lung. Further research is needed to disentangle the mechanisms behind this and determine whether senescence could be driving the age-associated ECM changes in the lung.

We used a unique approach to assess age-related ECM differences on levels and in specific lung compartment including the whole tissue transcript and protein analysis followed by extensive, immunohistochemical analyses in specific lung regions. Using immunohistochemical analysis, we observed an association of more COL1A1, COL6A2, COL14A1, and FBLN2 expression with higher age, while no association was found for COL6A1, LTBP4 and LUM expression. Sensitivity and specificity of the antibodies as well as differences in the epitope region recognized by the antibodies may explain the differences observed in results obtained from immunohistochemical analysis compared to the proteomic analysis. The differences observed in percentage area/distribution and mean intensity/expression of the positive stained age-associated ECM proteins provide different information about distribution and amount of protein present within the tissues.

Our study also had some limitation, namely the relatively small number of 9 control patients used for the proteomic analysis compared to the numbers of control patients, 270 and 64, used for transcriptomic analysis and immunohistochemical analysis, respectively. Additionally, our study is cross-sectional and therefore makes it difficult to infer any causal relationship.

As previously mentioned, the aging lung is characterized by structural and physiological alterations. As the ECM regulates different biomechanical properties of tissue and organs, and comprises key proteins responsible for lung stiffness, elasticity, and recoil. In addition, the different ECM proteins play a role in binding and release of specific cytokines and chemokines.

Therefore, the age-associated ECM differences that we showed with differences in several collagens as well as FBLN2, that co-localizes within the elastic fibre, are likely important contributors to structural and physiological changes in the aging lung.

In summary, our study revealed age-associated differences in the lung ECM from histologically normal lung from patients with normal lung function and no history of chronic lung disease. Higher COL6A2 expression with higher age was present in all lung compartments except in the bronchial epithelium, where it is not expressed. Most differences were observed in the parenchyma and vessel walls. These ECM differences may affect lung structure and physiology with aging and as such help in understanding the development and progression of chronic lung diseases. Identifying the mechanisms regulating the ECM deposition in the aging lung will lay a strong foundation for the identification of potential triggers for the development of age-associated chronic lung diseases.

## Funding

This study was partly supported by an Abel Tasman Talent Program Fellowship, in association with the Healthy Aging Alliance, provided by the University Medical Center Groningen and the Mayo Clinic, Rosalind Franklin Fellowship provided by the University of Groningen and the European Union, de Cock-Hadders Stichting grant provided by University Medical Center Groningen, NIH grants R01 HL088029 and R01 HL0142061. The proteomics analysis was supported by the Netherlands X-omics Initiative (NWO, project 184.034.019).

## Acknowledgements

The stainings of COL1A1, COL6A1, COL6A2, COL14A1, FBLN2, LTBP4 and LUM on lung tissue performed in this manuscript were conducted as part of the HOLLAND (HistopathOLogy of Lung Aging aNd COPD) project. The HOLLAND project was initiated and supervised by Corry-Anke Brandsma, Wim Timens, and Janette Burgess, technical support was provided by Marjan Reinders-Luinge, Anja Bakker and Theo Borghuis, and image analyses pipelines were developed by Theo Borghuis, Maunick Lefin Koloko Ngassie and Niek Bekker.

## Notes

### Competing Interest Statement

The authors have declared no competing interest.

### Summary of Updates

The supplementary file 1 has been updated. The previous version did not contain the references listed in the file. This has now been added at the end of the document.

## References

1. De Vries M, Faiz A, Woldhuis RR, Postma DS, De Jong TV, Sin DD, et al. Lung tissue gene-expression signature for the ageing lung in COPD. Thorax. 2018;73(7):609–17.

2. Zhou Y, Horowitz JC, Naba A, Ambalavanan N, Atabai K, Balestrini J, et al. Extracellular matrix in lung development, homeostasis and disease. Matrix Biology. 2018;73:77–104.

3. Meiners S, Eickelberg O, Königshoff M. Hallmarks of the ageing lung. European Respiratory Journal. 2015;45(3):807–27.

4. Cho SJ, Stout-Delgado HW. Aging and Lung Disease. Annual Review of Physiology. 2020;82(1):433–59.

5. Copley SJ, Wells AU, Hawtin KE, Gibson DJ, Hodson JM, Jacques AET, et al. Lung Morphology in the Elderly: Comparative CT Study of Subjects over 75 Years Old versus Those under 55 Years Old. Radiology. 2009;251(2):566–73.

6. Brandenberger C, Mühlfeld C. Mechanisms of lung aging. Cell and Tissue Research. 2017;367(3):469–80.

7. Huang K, Rabold R, Schofield B, Mitzner W, Tankersley CG. Age-dependent changes of airway and lung parenchyma in C57BL/6J mice. Journal of Applied Physiology. 2007;102(1):200–6.

8. Brandsma C-A, De Vries M, Costa R, Woldhuis RR, Königshoff M, Timens W. Lung ageing and COPD: is there a role for ageing in abnormal tissue repair? European Respiratory Review. 2017;26(146):170073.

9. Frantz C, Stewart KM, Weaver VM. The extracellular matrix at a glance. Journal of Cell Science. 2010;123(24):4195–200.

10. Starcher BC. Lung elastin and matrix. Chest. 2000;117(5 Suppl 1):229S–34S.

11. Chow RD, Majety M, Chen S. The aging transcriptome and cellular landscape of the human lung in relation to SARS-CoV-2. Nature Communications. 2021;12(1).

12. Burgstaller G, Oehrle B, Gerckens M, White ES, Schiller HB, Eickelberg O. The instructive extracellular matrix of the lung: basic composition and alterations in chronic lung disease. European Respiratory Journal. 2017;50(1):1601805.

13. Brandsma C-A, Guryev V, Timens W, Ciconelle A, Postma DS, Bischoff R, et al. Integrated proteogenomic approach identifying a protein signature of COPD and a new splice variant of SORBS1. Thorax. 2020;75(2):180–3.

14. Schindelin J, Arganda-Carreras I, Frise E, Kaynig V, Longair M, Pietzsch T, et al. Fiji: an open-source platform for biological-image analysis. Nature Methods. 2012;9(7):676–82.

15. Hunzelmann N, Nischt R, Brenneisen P, Eickert A, Krieg T. Increased deposition of fibulin-2 in solar elastosis and its colocalization with elastic fibres. British Journal of Dermatology. 2001;145(2):217–22.

16. Calhoun C, Shivshankar P, Saker M, Sloane LB, Livi CB, Sharp ZD, et al. Senescent Cells Contribute to the Physiological Remodeling of Aged Lungs. The Journals of Gerontology Series A: Biological Sciences and Medical Sciences. 2016;71(2):153–60.

17. Onursal C, Dick E, Angelidis I, Schiller HB, Staab-Weijnitz CA. Collagen Biosynthesis, Processing, and Maturation in Lung Ageing. Frontiers in Medicine. 2021;8.

18. Angelidis I, Simon LM, Fernandez IE, Strunz M, Mayr CH, Greiffo FR, et al. An atlas of the aging lung mapped by single cell transcriptomics and deep tissue proteomics. Nature Communications. 2019;10(1).

19. Cescon M, Gattazzo F, Chen P, Bonaldo P. Collagen VI at a glance. Journal of Cell Science. 2015;128(19):3525–31.

20. Mereness JA, Bhattacharya S, Wang Q, Ren Y, Pryhuber GS, Mariani TJ. Type VI collagen promotes lung epithelial cell spreading and wound-closure. PLOS ONE. 2018;13(12):e0209095.

21. Ansorge HL, Meng X, Zhang G, Veit G, Sun M, Klement JF, et al. Type XIV Collagen Regulates Fibrillogenesis. Journal of Biological Chemistry. 2009;284(13):8427–38.

22. Kobayashi N, Kostka G, Garbe JHO, Keene DR, Bächinger HP, Hanisch F-G, et al. A Comparative Analysis of the Fibulin Protein Family. Journal of Biological Chemistry. 2007;282(16):11805–16.

23. Ibrahim AM, Sabet S, El-Ghor AA, Kamel N, Anis SE, Morris JS, et al. Fibulin-2 is required for basement membrane integrity of mammary epithelium. Scientific Reports. 2018;8(1).

24. Patel MR, Weaver AM. Astrocyte-derived small extracellular vesicles promote synapse formation via fibulin-2-mediated TGF-ß signaling. Cell Reports. 2021;34(10):108829.

25. Khan SA, Dong H, Joyce J, Sasaki T, Chu M-L, Tsuda T. Fibulin-2 is essential for angiotensin II-induced myocardial fibrosis mediated by transforming growth factor (TGF)-ß. Laboratory Investigation. 2016;96(7):773–83.

26. Tsuda T, Wu J, Gao E, Joyce J, Markova D, Dong H, et al. Loss of fibulin-2 protects against progressive ventricular dysfunction after myocardial infarction. Journal of Molecular and Cellular Cardiology. 2012;52(1):273–82.

27. Sicard D, Haak AJ, Choi KM, Craig AR, Fredenburgh LE, Tschumperlin DJ. Aging and anatomical variations in lung tissue stiffness. American Journal of Physiology-Lung Cellular and Molecul ar Physiol ogy. 2018;314(6):L946–L55.

28. Lee S, Islam MN, Boostanpour K, Aran D, Jin G, Christenson S, et al. Molecular programs of fibrotic change in aging human lung. Nature Communications. 2021;12(1).

29. Wicher SA, Roos BB, Teske JJ, Fang YH, Pabelick C, Prakash YS. Aging increases senescence, calcium signaling, and extracellular matrix deposition in human airway smooth muscle. PLOS ONE. 2021;16(7):e0254710.

