## Supplementary file 1 for "Age-associated Differences in the Human Lung Extracellular Matrix"

#### **Immunohistochemical staining**

To localize the expression of the age-associated ECM proteins and to validate the transcriptomic and proteomic findings, immunohistochemical staining was performed in lung tissues derived from 64 control subjects. The lung tissues were embedded in paraffin and cut into 6 µM thick sections. These sections were deparaffinized and rehydrated; followed by antigen retrieval with 10 mM citrate buffer pH6 for COL1A1, FBLN2, LTBP4 and LUM stainings and 10 mM Tris/EDTA buffer pH9 for COL6A1, COL6A2 and COL14A1 stainings. Endogenous peroxidase activity was blocked by 0.3 % hydrogen peroxidase (H_2_O_2_), followed by overnight incubation at 4 ºC of the primary antibodies COL1A1 (monoclonal mouse, anti-human, ab88147, 1:400), COL6A1 (polyclonal rabbit, anti-human, NB120-6588, 1:3200), COL6A2 (monoclonal rabbit, anti-human, ab180855, 1:12500), COL14A1 (polyclonal rabbit, anti-human, HPA023781, 1:100), FBLN2 (polyclonal rabbit, anti-human, HPA001934, 1:400), LTBP4 (polyclonal rabbit, anti-human, ab222844, 1:400), and LUM (monoclonal rabbit, anti-human, ab168348, 1:12500) diluted in a 1% BSA-PBS. The sections were washed and incubated with the Horseradish peroxidase (HRP)-conjugated secondary antibodies diluted in 2% human serum + 1% BSA-PBS, i.e., polyclonal Goat Anti-Rabbit (P0488, Dako, Denmark) for COL6A1, COL6A2, COL14A1, FBLN2, LTBP4 and LUM and Rabbit Anti-Mouse (P0260, Dako, Denmark) for COL1A1 staining. Positive staining was visualized using 5 min incubation with Vector® NovaRED® Substrate (SK-4800, Vector Laboratories, Canada). Sections were counterstained with hematoxylin and scanned using the Hamamatsu NanoZoomer 2.0HT digital slide scanner (Hamamatsu Photonic K.K., Japan) at magnification of 40x. The digital images were viewed with Aperio ImageScope V.12.4.3 (Leica Biosystems, Germany).

#### **Image analysis**

Different compartments of the lung including the whole lung tissue, parenchyma, airway wall, airway epithelium and blood vessels were analyzed for the expression and distribution of ECM proteins. First, images containing the whole lung tissue, airways and blood vessels were extracted from the scans using Aperio ImageScope software V.12.4.3 (Leica Biosystems, Nussloch, Germany). Depending on how many airways and blood vessels were present in the tissue, up to ten airways or blood vessels were extracted. Next, Adobe Photoshop software (Adobe Inc. California, United States) was used to extract the specific areas for analysis (Figure 1B), i.e. parenchyma excluding airway and vessels, airway wall from basement membrane to alveolar attachments, bronchial epithelial layer from basement membrane to ciliary layer and blood vessel walls from endothelium to alveolar attachment except for LUM (from tunica media to endothelium), since LUM was localized in that specific region. Artifacts including carbon pigments, folded tissue, red blood cells and mucus plugs were removed. Fiji/ImageJ software (1) was used to quantify the intensity and area of positive staining (Figure 1C). All image files were separated into blue (hematoxylin-image), and red (NovaRed-image) pixels using colour deconvolution plugin by Landini and colleagues (2). To determine the correct optical density vectors for the red-green-blue (RGB) channel of Hematoxylin and NovaRed, we followed the protocol as previously described by Ruifrok and colleagues (3). A macro was developed to receive automated numbers for each pixel in the different images. To calculate the total amount of tissue, images were converted to 8-bit grey scale. Total number of pixels representing total tissue area versus positively stained tissue area were identified using the threshold feature of Fiji/ImageJ software. The percentage of positive area and the mean intensity were calculated using the following formulas, where the percentage positively stained tissue (Area %) is calculated by dividing positive stained NovaRed pixels by the total amount of grey scale pixels representing total tissue area. The protocol described previously by Nguyen was used for the calculation of the mean intensity which represent the which in proportional to the expression of protein (4). The pixel intensities of separated NovaRed images range from 0 to 255 in Fiji/ImageJ software, where the value 0 represents the darkest shade of the color, while 255 represents the lightest shade of color in the image. Hereby, it should be remembered that the darker the positive NovaRed pixels are, the smaller their intensities are. Thus, the reciprocal intensity is calculated by subtracting the value obtained by dividing the total number of NovaRed pixels by the area of positive NovaRed staining from 255. The reciprocal intensity is defined as mean intensity and is proportional to the amount of positive NovaRed pixels present in the analyzed image. With this method, we determined the expression level of proteins we stained in lung tissues. The analysis of data and calculations were performed using R software V.4.0.0 (Boston, Massachusetts, USA).

$$Area (\%)=\frac{Number of pixels positive for NovaRed}{number of pixels in total tissue}*100$$

$$Mean intensity=255-\frac{Sum of intensities of pixels positive NovaRed}{Total number of pixels positive for NovaRed}$$

| **Table S1** | **Subject characteristics transcriptomics cohort** | | |
| --- | --- | --- | --- |
|  | **Groningen** | **Vancouver** | **Quebec** |
| Number | 45 | 90 | 135 |
| Age, years (range) | 60 (37-76) | 62 (40-80) | 62 (41-80) |
| Male/female, N | 20/25 | 49/40 | 76/59 |
| Smoking, N |  |  |  |
| Ex | 25 | 59 | 113 |
| Current | 20 | 31 | 22 |
| Pack years, N | 35 (21-41) | 37.75 (24-49.9) | 38.35 (25-46) |

| **Table S2** | **Subject characteristics proteomics cohort** | |  |  |  |  |
| --- | --- | --- | --- | --- | --- | --- |
|  |  | **Age** | **Sex (m/f)** | **Packyears** | **FEV_1_%pred** | **FEV_1_/FVC ratio** |
| Non-COPD control (n=9) | | 66 (6) ^#^ | 4/5 | 34 (17) * | 94 (10) | 76 (4) |
| mean (stdev) |  |  |  |  |  |  |
| * 2 controls had missing info for packyears |  |  |  |  |  |  |
| ^#^ p<0.05 control versus COPD |  |  |  |  |  |  |

#### **References**

1.	**Schindelin J, Arganda-Carreras I, Frise E, Kaynig V, Longair M, Pietzsch T, Preibisch S, Rueden C, Saalfeld S, Schmid B, Tinevez J-Y, White DJ, Hartenstein V, Eliceiri K, Tomancak P, and Cardona A**. Fiji: an open-source platform for biological-image analysis. *Nat Methods* 9: 676-682, 2012.

2.	**Landini G, Martinelli G, and Piccinini F**. Colour deconvolution: stain unmixing in histological imaging. *Bioinformatics* 37: 1485-1487, 2021.

3.	**Ruifrok AC, and Johnston DA**. Quantification of histochemical staining by color deconvolution. *Anal Quant Cytol Histol* 23: 291-299, 2001.

4.	**Nguyen D**. Quantifying chromogen intensity in immunohistochemistry via reciprocal intensity. *Protoc Exch* 2013.
